# Regulation of B cell migration to the Bursa of Fabricius by CCR7 and cell adhesion molecules in chicken embryonic development

**DOI:** 10.1101/2024.12.04.626876

**Authors:** Milena Brunner, Catarina L. C. T. Cavaleiro, Tom V. L. Berghof, Christine Wurmser, Daniel Elleder, Theresa von Heyl, Benjamin Schusser

## Abstract

The development of functional B lymphocytes during chicken embryogenesis relies on a series of tightly regulated processes. Precursor B cells migrate from the spleen via the blood to the bursa of Fabricius, where they colonize the bursal follicles to undergo further maturation and differentiation. To better understand the molecular mechanisms underlying early B cell migration in the chicken embryo, transcriptome analysis of B cells isolated from the spleen, blood, and bursa at embryonic days (ED) 12, ED14, and ED16 was performed. These findings suggest that sphingosine-1-phosphate (S1P) and its receptors regulate B cell presence in the bloodstream, while CCR7 and CXCR4 guide B cells to the bursa. Additionally, integrins and cell adhesion molecules, such as PECAM1, appear to facilitate transendothelial migration into the bursal mesenchyme. This study highlights a coordinated interplay between chemokines, integrins and cell adhesion molecules involved in B cell recruitment and colonization of the bursa microenvironment. These findings enhance our understanding of early B cell migration and shed light on the mechanisms governing B cell trafficking during chicken embryonic development.

## Introduction

Developing highly specialized B lymphocytes in chickens involves a series of precise developmental changes within the embryo. Despite significant progress, several aspects of chicken B cell development and the mechanisms to build up a robust humoral immune response are still not fully investigated. During chicken embryonic development, precursor B cells arise from hematopoietic cells in the dorsal aorta and subsequently populate the spleen [1, 2]. Starting at embryonic day (ED) 10, these immature B cells exit the spleen, enter the bloodstream, and colonize the bursa of Fabricius, a specialized avian organ required for B cell development [3–5]. Chicken B cells must colonize the bursa to undergo several developmental changes and progress into fully mature B cells [6–8]. Inside the bursal follicles, the B cell receptor (BCR) repertoire diversification occurs through V(D)J gene rearrangement followed by somatic gene conversion [9, 10]. Around hatching, these newly diversified B cells leave the bursa and populate secondary peripheral lymphoid tissues to participate in immune responses [11].

The colonization of the bursal microenvironment by B cell progenitors is a crucial phase of B cell development and occurs in two stages: migration into the bursal mesenchyme and subsequent colonization of the lymphoid follicles. Despite compelling evidence that B cell progenitors are already committed to the B cell lineage [1], more work needs to be done to fully understand the underlying mechanisms. It has been shown that V(D)J gene rearrangement can occur in the pre-bursal stage [12, 13], and B cells leaving the bursa depend on surface immunoglobulin (sIg) expression [14, 15]. Moreover, recent studies demonstrated, that the interaction of CXCR4/CXCL12 seems to play a crucial role in chicken B cell migration to and out of the bursa [16]. CXCR4^+^-expressing B cells migrate towards CXCL12 produced in the bursal anlage and follicle buds, indicating a CXCR4/CXCL12-mediated mechanism of B cell migration [16, 17]. However, B cells lacking the BCR stopped migrating towards CXCL12, suggesting that while the CXCR4/CXCL12 axis is crucial for B cell migration, the presence of surface BCR is necessary to trigger the signaling cascade [17]. However, sIg expression is not necessary for precursor B cells to enter the bursal mesenchyme, as BCR^neg^ B cells can colonize the bursa normally [15, 18]. Based on these findings, we propose that the CXCR4/CXCL12 axis is not the exclusive driver and that additional factors are needed for directed migration to and exclusive B cell colonization of the bursa during chicken embryonic development.

Earlier studies examining the recruitment of pre-bursal B cells to the bursa have identified distinct expression patterns in cell surface glycosylation during lymphoid cell development, where pre-bursal B cells express sialyl-Lewis(x) (CD15s) before switching to Lewis(x) (CD15) during gene conversion. CD15s is suggested to facilitate adhesion through selectin binding [19, 20]. The loss of CD15s around ED15 correlates with the loss of the colonization potential of pre-bursal B cells [21]. In rabbits, the interaction between selectin L (CD62L) and PNAd, along with CCR7 and the integrins LFA1 and α4ß1, seem to be mandatory for B cell migration and appendix colonization, indicating a mechanism involving specific glycosylation patterns and adhesion molecules [22, 23]. Additionally, sphingosine-1-phosphate (S1P) is a phospholipid produced by sphingosine kinases (e.g., SPHK1), acting as a chemoattractant for hematopoietic stem cells (HSCs) by binding to its receptors (e.g., S1P1). High S1P levels in the blood and low levels in the tissue create a gradient that is used by HSCs and lymphocytes for tissue egress [24, 25]. The egress of HSCs from the bone marrow in mice relies on S1P/S1P1 interaction, highlighting a potential contribution of CXCR4 antagonist-mediated mobilization of the HSCs [26]. In the adult hen, S1P1 appears to regulate immune cell infiltration into ovarian cancers, and chicken embryo amacrine cells have been reported to express S1P [27, 28]. However, the participation of sphingosines in the context of B cell migration in the chicken has not been studied so far.

Our research aims to investigate the mechanism driving early B cell migration and identify the signals involved in this process. We hypothesize that early B cell migration is mediated through signals located on the cell surface or receptors in the plasma membrane that direct B cells to their target tissue. To examine this, we performed RNA-sequencing to profile the B cell transcriptome isolated from the spleen, blood, and bursa at ED12, ED14, and ED16. This approach enabled us to identify differentially expressed genes (DEGs) within the integrins, cell adhesion molecules, and the chemokine receptor gene families, providing valuable insights into the molecular mechanisms that drive early B cell migration in the chicken embryo.

## Results

### Temporal changes in B cell populations and transcriptome during embryonic B cell development

This study aimed to identify the signals guiding precursor B cells to the bursa and facilitate their transmigration from the blood to the bursal mesenchyme during chicken embryonic B cell development. B cell samples were collected from the spleen, blood, and bursa, allowing for the analysis of the B cell transcriptome as they migrate from the embryonic spleen through the bloodstream to the bursa of Fabricius on ED12, ED14, and ED16. Lymphocytes were isolated, and flow-cytometry-based cell sorting was used to isolate the B cells (Fig. 1A and S1). The percentages of B cells within living peripheral blood mononuclear cells (PBMCs) varied over time between the different tissues. On ED12, a higher percentage of B cells was found in the spleen (6%) compared to the blood (0.6%) and the bursa (1.5%). Slightly higher percentages of B cells were present in the spleen on ED14 (8%), while the lowest percentages were observed on ED16 (2.9%). Over time, B cell percentages increased in the bursa significantly from ED12 (1.5%) to ED14 (13%, *p*=0.009), reaching the highest percentage at ED16 (32%, *p*<0.001). In contrast, no changes in B cell percentages were observed in the blood (0.4-0.6%) (Fig. 1B). Principal Component analysis (PCA) revealed that B cell samples from the spleen and blood formed distinct but closely positioned clusters, indicating their transcriptional similarity. In contrast, separate clusters for each time point were observed for bursal B cells. At ED14 bursal B cells co-localized with those from the spleen and blood, while bursal B cells on ED12 and ED16 formed separate and isolated clusters, highlighting their transcriptomic distinctiveness (Fig. 1C).

**Figure 1:**
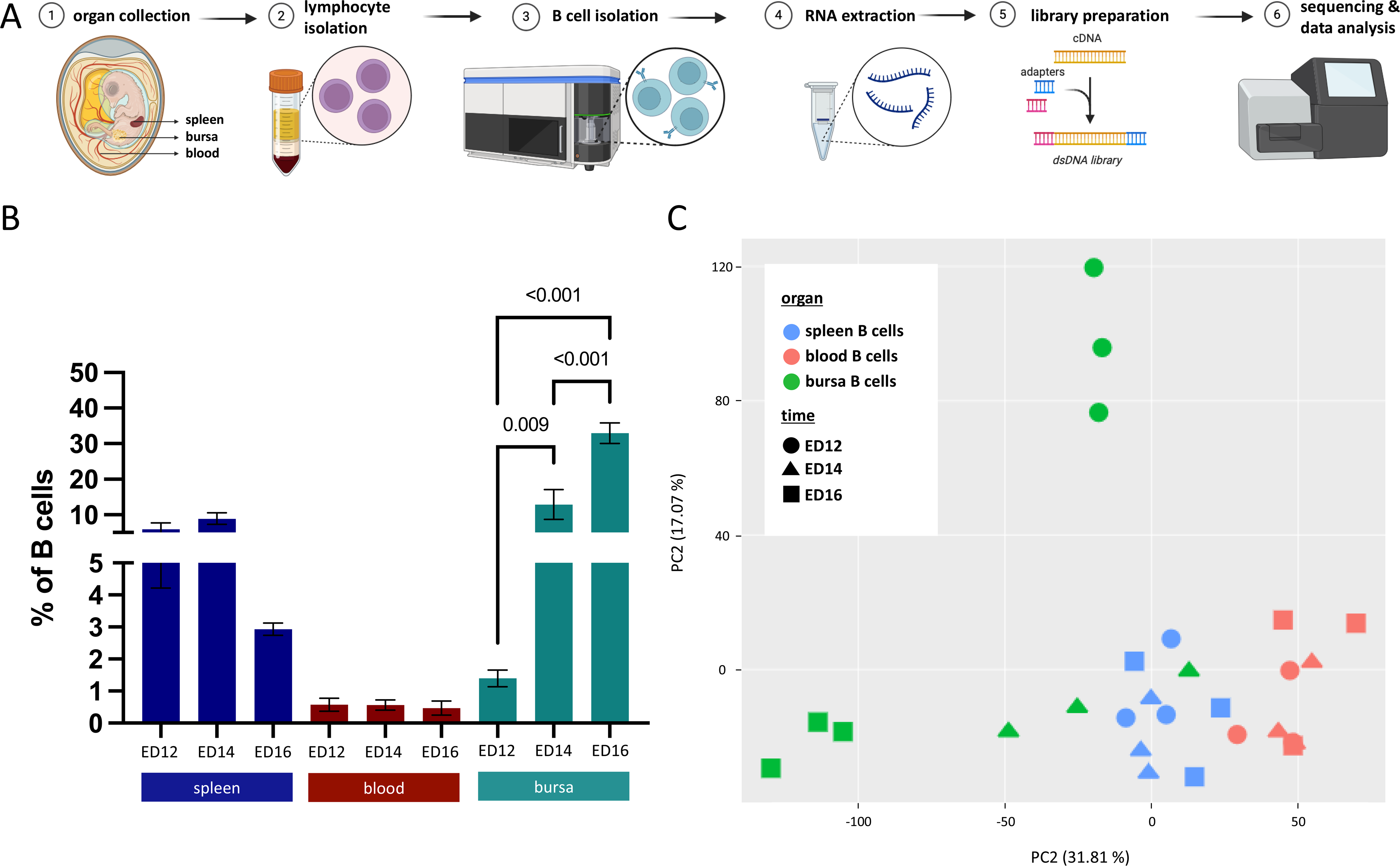
RNA-sequencing analysis of B cell migration in the chicken embryo. (A) Schematic representation of the workflow from organ collection, separation of lymphocytes, isolation of B cells by flow-cytometry-based cell sorting, RNA extraction, library preparation, NGS, and data analysis. (B) Flow-cytometry analysis of B cells (Bu1-FITC^+^) within single live cells (LD-eFluor780^-^) in percentages from the spleen (blue), blood (red), and bursa (green) extracted on ED12 (left), ED14 (middle) and ED16 (right). Bars represent the mean ± SEM and p-values (one-way ANOVA) are indicated in the graph. (C) Principal-Component-Analysis (PCA) from splenic (blue), blood (red), and bursal (green) B cell samples collected at ED12 (circle), ED14 (triangle), and ED16 (square). Gene expression data was normalized with MRN, and batch-corrected with Combat. (B+C) Data are cumulative from 3 experiments (*n*=3 per B cell sample and time point).

### The bursa exhibits the highest numbers of DEGs, independent of the time point

To gain insight into the transcriptome of migrating B cells within the chicken embryo, pairwise differential expression analysis between the B cell populations of each tissue and time point revealed distinct gene expression patterns in each tissue, particularly in the bursa (Fig. 2). In contrast, most genes exhibit similar expression patterns in the spleen and blood that remained consistent over time. (Fig. 2A). Up- and downregulation of DEGs were analyzed in B cells isolated from the different tissues across time points (Fig. 2B and S2). The highest number of DEGs were detected for blood vs. bursa on ED12 (1045 up 1071 down) and ED16 (1624 up 1885 down), compared to the other tissue comparisons. Additionally, the highest number of DEGs in the spleen vs. bursa were observed at ED16 (1363 up 1597 down), whereas the lowest was seen on ED14 (77 up 45 down). In contrast, the number of DEGs in the spleen vs. blood comparison showed minor variation over time (ED12: 228 up 269 down; ED14: 296 up 336 down; ED16: 419 up 134 down). In the time point comparison, the bursa displayed the highest number of DEGs, and most were detected for ED12 vs. ED16 (1432 up 1805 down), while the fewest on ED12 vs. ED14 (345 up 654 down). In the spleen, the highest DEG count occurred at ED14 vs. ED16 (27 up 110 down), whereas low numbers were observed for both ED12 vs. ED14 (18 up 7 down) and ED12 vs. ED16 (13 up 5 down). No DEGs were found in the blood at ED12 vs. ED14, and only few were observed at ED14 vs. ED16 (10 up 5 down), as well as for ED12 vs. ED16 (3 up 5 down) (Fig. 2C).

**Figure 2:**
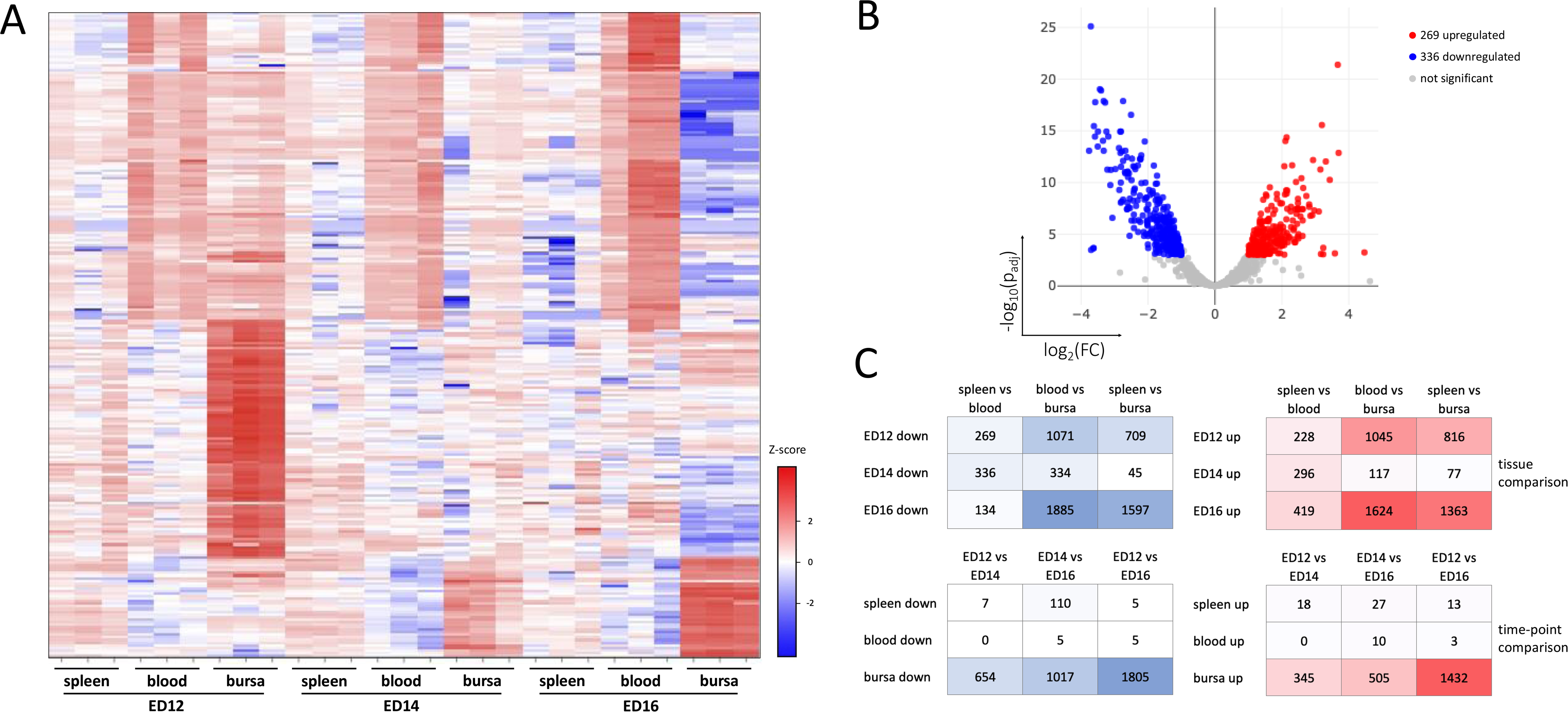
Analysis of differentially expressed genes (DEGs) across tissues and time points. (A) Heatmap of the 250 most-varied DEGs with the highest p_adjust_/fold-change between the tissues and time points. (B) Representative Volcano plot to display the variation in gene expression between spleen vs. blood on ED14. The log_2_(FC) indicates the mean fold-change (FC) in expression levels for each gene, and each dot represents one gene. DEGs identified in pairwise comparisons between the tissues and time points are defined as p_adjust_ < 0.001, FC > |2|. DEGs with higher expression (i.e., higher in the spleen than in blood) are represented by red dots, while DEGs with lower expression (i.e., lower in the spleen than in blood) are denoted by blue dots. Genes with no substantial change in expression are indicated as gray dots. (C) The table shows the number of up- (right) and downregulated (left) DEGs between condition 1 and condition 2 (conditions being tissues (top) or time points (bottom)). The intensity of the color (red = higher in condition 1 than condition 2 and blue = lower in condition 1 than condition 2) corresponds to the number of DEGs.

### Functional classification of the DEGs

Gene Ontology (GO) analysis was used to examine the putative functions of the DEGs, categorizing them into biological process (BP), molecular function (MF), and protein class (PC). This functional classification provided insights into the processes predominantly affected by the DEGs across the tissue comparisons. The percentages of genes involved in various processes within each category remained consistent, regardless of the number of DEGs identified across the comparisons. DEGs from the tissue comparison were mainly associated with cellular processes (32.7%) within the BP category, followed by biological regulation (22.1%), metabolic process (14.1%), and response to stimulus (10.2%). B cell samples exhibited enrichment in MF categories for binding (34.6%) and catalytic activity (34.3%), while within the PC category, enrichment for metabolite interconversion enzyme (19.8%), protein modifying enzyme (12.3%), and protein-binding activity modulator (10.3%) was observed. DEGs associated with BP term locomotion and cell adhesion molecules within the PC category were particularly relevant to this study: 0.6% of the DEGs were linked to locomotion, and 1% to cell adhesion molecules (Fig. 3A).

**Figure 3:**
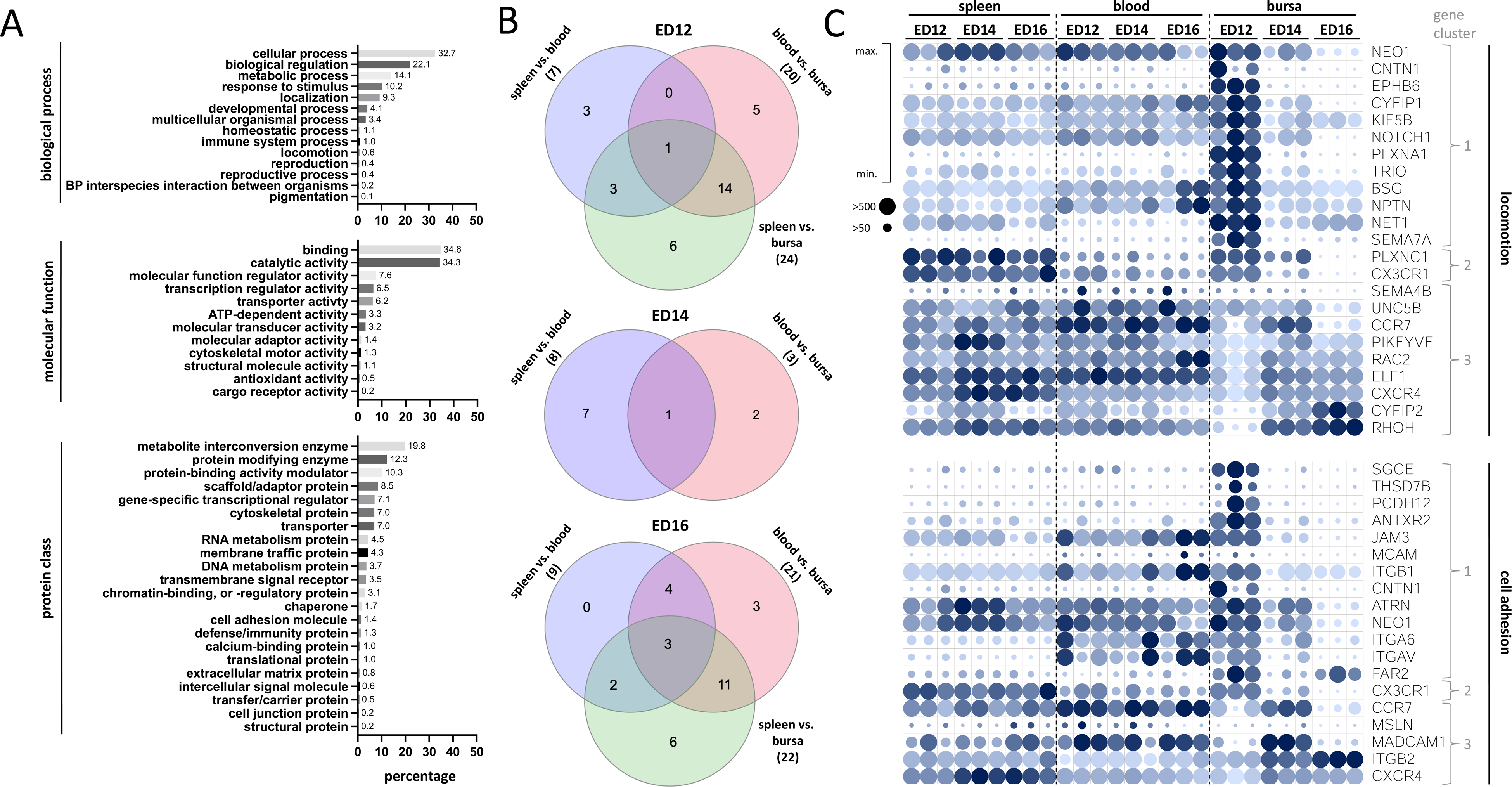
DEGs associated with GO terms locomotion or cell adhesion. (A) Gene Ontology (GO) analysis of DEGs (p_adjust_ < 0.001, fold-change > |2|). The DEGs were examined for their enrichment in the three GO ontologies: biological process (top), molecular function (middle), and protein class (bottom). Above each individual bar is the percentage of genes enriched in the corresponding GO terms. (B) Venn diagram of shared DEGs between the tissues spleen vs. blood (blue), spleen vs. bursa (green), blood vs. bursa (red) at ED12 (top), ED14 (middle) and ED16 (bottom). (C) Gene expression profile of DEGs associated with GO-terms locomotion (top) or cell adhesion molecules (bottom) between the tissues over time. The Pearson correlation coefficient was used for the nearest neighbor analysis, and genes were clustered based on similar expression profiles. Coloring of the circles is based on the minimal (light blue) and the maximal (dark blue) value of each gene, and the size of the circles determines the overall expression value, small circles (<50) and big circles (>500) gene counts.

### Dynamic expression of genes related to locomotion and cell adhesion molecules

For the subsequent analysis, we focused only on genes identified in locomotion and cell adhesion molecules in at least one of the tissue comparisons (n=37). Venn diagrams were used to identify genes uniquely expressed in one of the three tissues as well as genes shared across two or all tissues at different time points (Fig. 3B and Tab. S1). Gene clustering based on expression profiles across the tissues over time aimed to further investigate the expression patterns of selected DEGs. Nearest-neighbor analysis and Pearson correlation effectively grouped locomotion-related genes and cell adhesion molecules into three distinct clusters. Cluster 1 (*NEO1, CTN1, EPHB6, KIF5B, NOTCH1, PLXNA1, TRIO, BSG, NPTN, NET1*, *SEMA7A, SGCE, THSD7B, PCDH12, ANTXR2, JAM3, MCAM, ITGB1, ATRN, ITGA6, ITGAV and FAR2)* exhibit relatively high gene expression in the bursa at ED12, with moderate to low expression in other tissues and time points. Cluster 2 genes (*PLXNC1* and *CX3CR1*) displayed high expression in the spleen, decreased expression in the bursa over time, and low expression in the blood. Cluster 3 genes (*SEMA4B, UNC5B, CCR7, PIKFYVE, RAC2, ELF1, CXCR4, CYFIP2, RHOH, MSLN, MADCAM1* and *ITGB2)* displayed consistently high expression in either the spleen or the blood, and dynamic expression patterns in the bursa over time, showing the highest expression on ED12 and lowest on ED16 or vice versa, except for *CCR7* and *MADCAM1*, which peaked at ED14 (Fig. 3C).

### Temporal expression of migration-related genes revealed tissue-specific regulation

To further investigate potential regulators of migration, we examined the temporal expression profiles of gene families related to migration: integrins, cell adhesion molecules (CAMs), sphingosine and chemokine receptors (Fig. 4A and B). In the spleen, expression levels of integrins, CAMs and sphingosines did not change over time, except for a significant increase in *S1PR1* expression at ED14 (*p*=0.006). In the blood, an upregulation in integrin expression, particularly *ITGA6, ITGB1, ITGB1BP1, ITGB2* and *ITGAV* was detected at ED16. In the bursa, *ITGA6*, *ITGB1*, *ITGAV* and *ITGB4* (*p*<0.001) expression decreased over time, whereas *ITGB2* expression increased significantly (*p*<0.001). *PECAM1* exhibited very high expression levels, exceeding 20.000 gene counts in both the spleen and blood, in contrast to other CAMs. However, *PECAM1* expression significantly dropped 50-fold from ED14 to ED16 (*p*=0.027) in the bursa. *JAM3* expression increased in the blood from ED14 to ED16 but decreased significantly in the bursa from ED12 to ED14 (*p*=0.012) and from ED12 to ED16 (*p*=0.003). In the bursa, *MADCAM1* showed a 4-fold higher expression on ED14 than at ED12 and ED16. A decline in *S1PR1* expression in the blood was observed over time. Both *S1PR1* and *S1PR4* exhibited elevated expression at ED14 in the bursa, while a significant decrease in *S1PR1* expression was detected from ED14 to ED16 (*p*=0.002) (Fig. 4A). Additionally, the gene expression profile of enzymes involved in the synthesis of (sialyl)-Lewis(x) were analyzed. In the blood, an increase in *ST3GAL6* expression was observed between ED14 and ED16. The highest *ST3GAL5* expression was detected at ED14 in the spleen, while its expression in the blood decreased over time (Fig. S3A).

**Figure 4:**
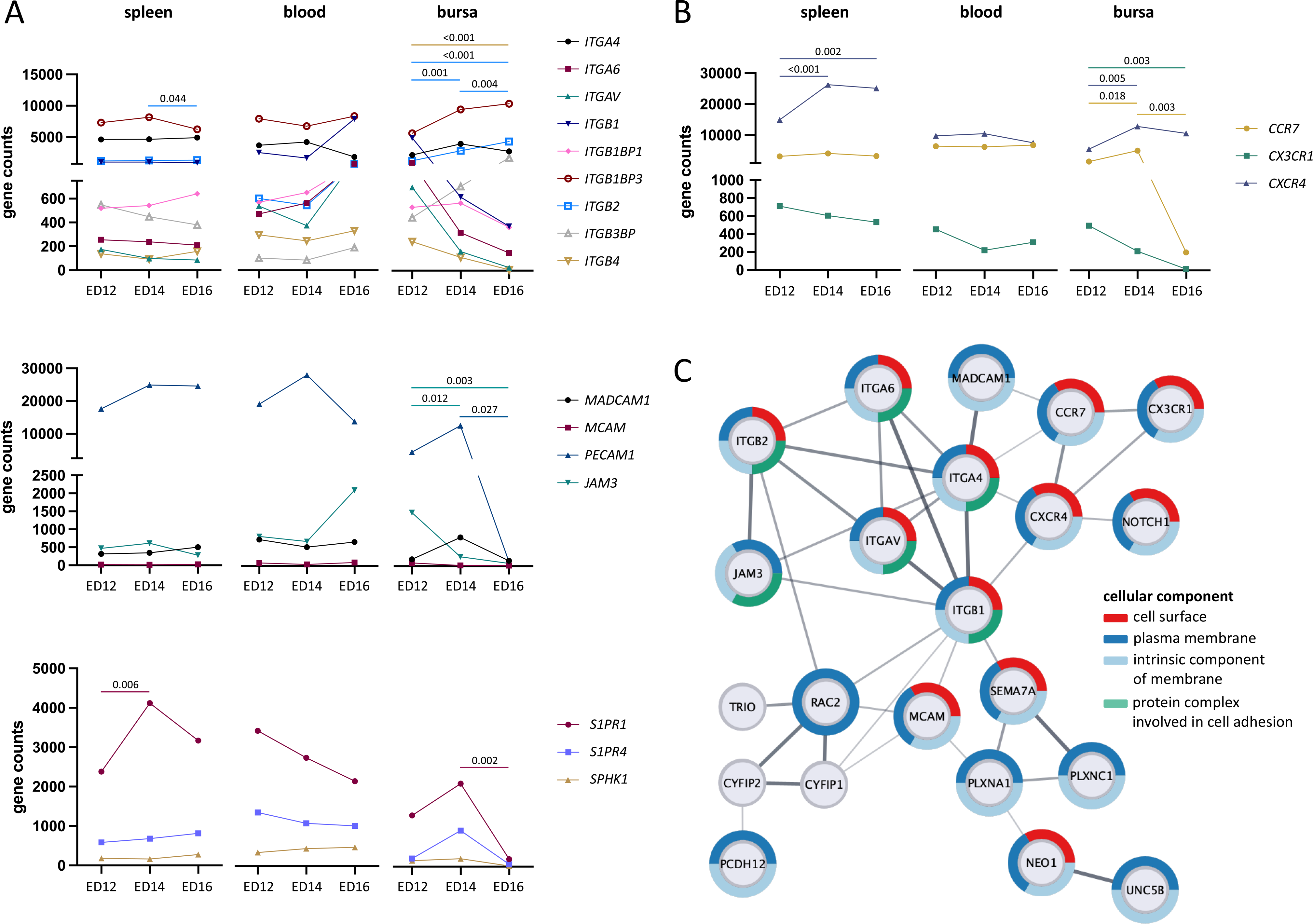
Expression profiles of gene sets related to lymphocyte migration exhibit different expression pattern in the tissues over time. (A+B) Expression profiles of the gene set integrins, cell adhesion molecules, sphingosines, and chemokine receptors over time (ED12, ED14, and ED16) in the spleen (left), blood (middle), and bursa (right). Each symbol represents an individual gene, with cumulative data from 3 experiments (*n*=3 per B cell sample and time point) shown as mean value, and statistical significance (one-way ANOVA) indicated by p-values in the graph. (C) PPI network from the DEGs associated with locomotion and cell adhesion are shown as nodes and presented as String network. Two nodes are connected if there are known interactions, and the thickness of the edge is based on evidence. DEGs were found significantly enriched (p-value < 0.05) in the category cellular compartment (CC) cell surface, plasma membrane, intrinsic component of membrane and protein complex involved in cell adhesion. Coloring and proportion around the nodes represent the enrichment within the top four CC categories cell surface (red), plasma membrane (dark-blue), intrinsic component of membrane (light-blue), and protein complex involved in cell adhesion (green).

### Time-dependent expression pattern of chemokine receptors in bursal B cells

Within the chemokine receptors, *CCR7* maintained consistently high expression levels in the blood and 1500 gene counts lower in the spleen. *CCR7* expression in the bursa significantly decreased between ED14 and ED16 (*p*=0.003), dropping from over 5000 gene counts to just 200. *CX3CR1* was more abundant in the spleen than in other tissues, while *CX3CR*1 expression significantly declined in the bursa over time (*p*=0.003). Interestingly, *CCR7* and *CX3CR1* expression levels in the spleen and blood did not show substantial variations between the time points. A significant increase in *CXCR4* expression from ED12 to ED14 (*p*<0.001) and ED16 (*p*=0.002) is observed in the spleen. Moreover, *CXCR4* expression increased significantly from ED12 to ED14 (*p*=0.005) in the bursa but decreased from ED14 to ED16 (Fig. 4B).

### Subcellular localization of locomotion and cell adhesion molecules

The putative signals involved in guiding B cells to the target tissue are supposedly located at the cell surface. Therefore, functional enrichment analysis was conducted and the subcellular localization of the DEGs was addressed by examining the top four GO terms in the cellular component (CC) category. The integrins and chemokine receptors, along with MCAM, SEMA7A, NEO1, and NOTCH1, were predominantly localized at the cell surface, plasma membrane, and as intrinsic component of the membrane. In contrast, MADCAM1, PLXNC1, PLXNA1, UNC5B, and PCDH12 were exclusively associated with the plasma membrane. The integrins and JAM3 were also linked to the CC term protein complex involved in cell adhesion. TRIO, CYFIP1, and CYFIP2 were localized to other cellular compartments (Fig. 4C and Tab. S2).

### Integrins as dominant interaction partners

Protein-protein interaction (PPI) network analysis revealed that 22 of the 37 DEGs, which encoded proteins associated with locomotion and cell adhesion, were likely to interact with each other, as shown in the STRING network. Notably, integrins, chemokine receptors (CCR7, CXCXR4, and CX3CR1), and CAMs (JAM3, MCAM, and MADCAM1) had multiple interaction partners, with ITGB1 being the most connected protein (n=9) within the network (Fig. 4C).

### Cell surface molecules are predominantly linked to extracellular matrix (ECM) organization and transendothelial migration pathways

Next, we investigated the DEGs not linked to a specific GO term but those located on the cell surface. Therefore, the subcellular localization of DEGs was analyzed from the comparisons of blood or spleen vs. bursa. From 1364 DEGs identified in the blood vs. bursa comparison, 38 were associated with the cell surface. Similarly, 52 out of 774 DEGs from the spleen vs. bursa comparison were located at the cell surface (Fig. 5A and Tab. S3). Functional enrichment analysis of these cell surface DEGs revealed significant enrichment (*p*<0.05) of KEGG pathways for cell adhesion molecules, leukocyte transendothelial migration, regulation of the cytoskeleton, and ECM-receptor interaction. Reactome pathways were found to be significantly enriched (*p*<0.05) for ECM-related pathways such as ECM degradation and organization, cell surface interactions at the vascular wall, and integrin cell surface interaction. Moreover, pathways related to immune responses, like cytokine signaling in the immune system and signaling by interleukins, were also found to be significantly enriched (Fig. 5B). The majority of cell surface DEGs identified from the blood or spleen vs. bursa comparisons were present in the KEGG pathway cell adhesion molecules involved in leukocyte transendothelial migration (Fig. 5C).

**Figure 5:**
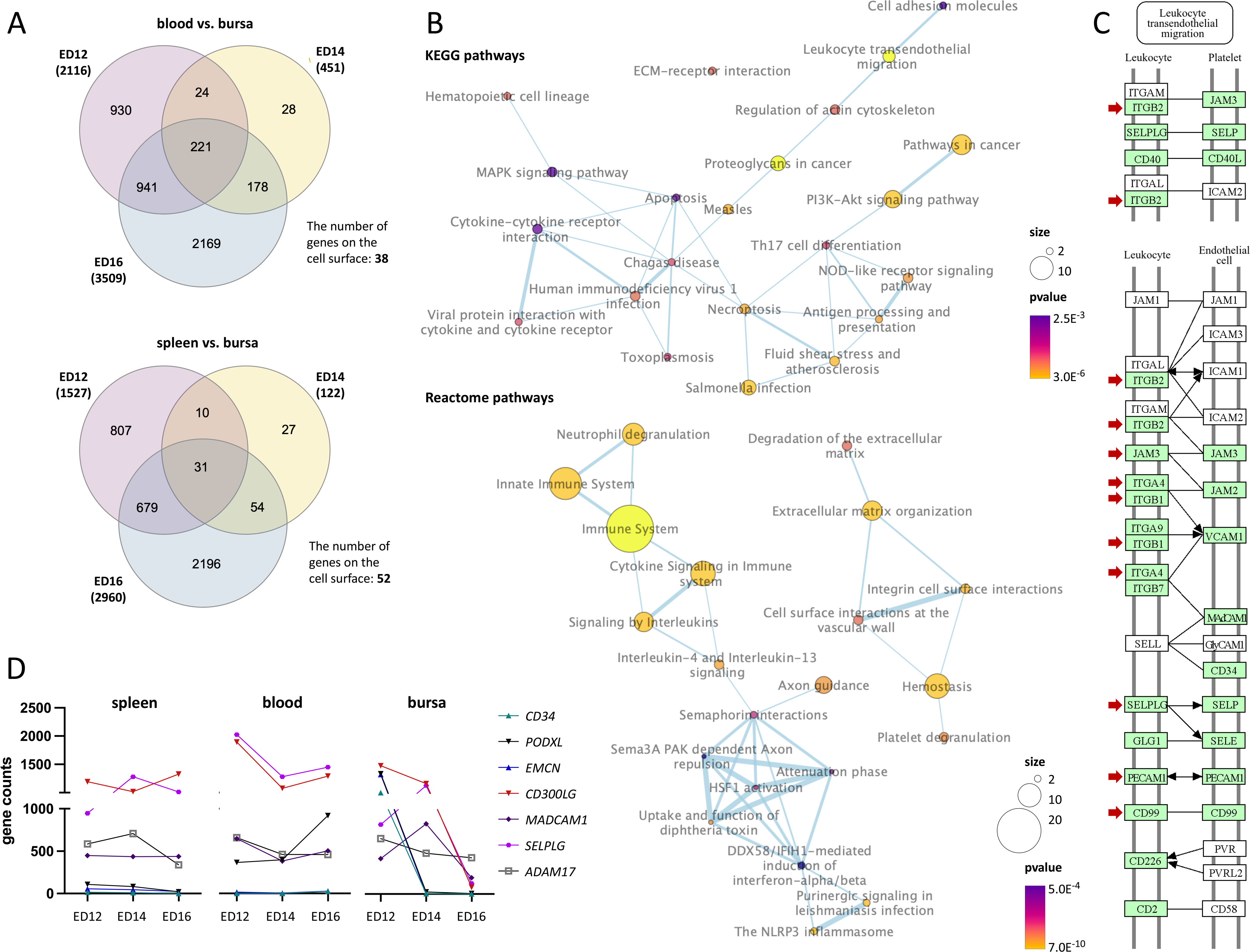
Cell surface DEGs are enriched in pathways linked to ECM and leukocyte transendothelial migration. (A) Venn diagrams show the number of shared DEGs from the blood vs. bursa (top) and spleen vs. bursa (bottom) comparisons on ED12 (red), ED14 (yellow), and ED16 (blue). DEGs shared across at least two time points were analyzed for their presence in the gene set ‘cell surfac’. (B) Significantly (p-value <0.05) enriched KEGG (top) and Reactome (bottom) pathways identified from the cell surface DEGs in the blood vs. bursa comparison. The number of genes in each pathway is reflected by circle size and the color gradient (yellow to violet) indicates significance (p-value). Thicker connecting lines between nodes represent more shared genes between pathways. (C) The KEGG pathway map for chicken cell adhesion molecules highlights their involvement in leukocyte transendothelial migration. Genes present in chickens are shown in green, while genes absent in chickens are displayed in white. Genes identified as DEGs in any of the tissue or time point comparisons were marked with a red arrow. (D) Gene expression profiles of known interactors or ligands of SELL over time in the spleen (left), blood (middle), and bursa (right).

### Dynamic expression of SELL interactors and phylogenetic analysis of selectins

In addition, the gene expression profiles of potential SELL interactors were analyzed in the different tissues over time. *SELPLG* expression in the spleen increased 1.5-fold from ED12 to ED14. In the blood, *CD300LG* and *SELPLG* expression decreased between ED12 and ED14, while *PODXL* expression increased from ED14 to ED16. In the bursa, *EMCN*, *CD34*, and *PODXL* expression decreased from over 1500 gene counts to almost zero. Additionally, an increase in *SELPLG* expression is observed from ED12 to ED14, which dropped from ED14 to ED16. A comparable decrease in expression was detected for *CD300LG* in the bursa (Fig. 5D). The highest expression of *SELE* and *SELP* was detected in the blood compared to the spleen and bursa, with the lowest expression levels in the blood at ED14 and the highest at ED16 (Fig. S3B). Moreover, the presence of SELL in chickens was analyzed by constructing a phylogenetic tree. This analysis revealed that all avian sequences labeled as L in GenBank clustered with human E, except for the chicken sequence, which may indicate a potential mislabeling of avian L sequences. Comparison of the different selectin sequences with each other, identified three distinct clusters, with no avian sequence grouping with the L cluster (Fig. S5).

## Discussion

Understanding the developmental changes that contribute to the maturation of B cell precursors in the chicken embryo is crucial. Originating from the dorsal aorta, the migration of immature B cells from the spleen to the bursa and their subsequent colonization of the bursal mesenchyme and follicles is a mandatory process during chicken B cell development in the embryo [2–5]. Even though several studies have been conducted to investigate the recruitment of precursor B cells to the bursa using diverse *in vitro* and *in vivo* models and techniques [15–17], the underlying mechanisms are still not fully identified.

This study employed a novel approach by analyzing the B cell transcriptome across all relevant tissues collected from the same embryos at different time points to advance understanding of the signals guiding B cell recruitment and entry into the bursal mesenchyme. B cells were isolated at an early stage from ED12 to ED16 and at each time point, the spleen, blood, and bursa were collected from the same embryos. This unique approach enabled continuous tracking of B cell migration across tissues and developmental stages, providing new insights into the dynamics of B cell trafficking in chicken embryos. Previous studies showed that B cell migration from the spleen to the bursa occurs in one wave, which takes place between ED8 and ED15 during embryonic development [5]. In consistency, B cell percentages decreased in the spleen while accumulating in the bursa over time, whereas continuously low percentages were detected in the blood.

Interestingly, bursal B cells seem to undergo several developmental changes, as indicated by a greater number of DEGs and distinct clusters in the PCA blot compared to B cells from the blood and spleen. This probably reflects the rapid transitions through several developmental stages B cells undergo as they mature. The transcriptome of bursal B cells changed severely over time, in contrast to the spleen and blood, where only small changes were observed. Only a small number of B cells have successfully entered the bursal mesenchyme at ED12, while at ED14, most B cells are still moving towards the bursa (Fig. 1B). By ED16, most B cells successfully entered the bursa and presumably colonized the follicle buds, to continue their maturation [16, 29].

It was shown recently that the interaction of CXCR4 and its ligand CXCL12 is essential for migrating B cells during chicken embryonic development, where CXCR4^+^ B cells migrate towards CXCL12 expressed by stromal or bursal secretory dendritic cells in the bursa [16, 17]. In our transcriptome analysis, *CXCR4* was identified as differentially expressed, with continuous high expression among the different tissues, which seems to be independent of the time point confirming previous findings. Since the contribution of CXCR4/CXCL12 in chicken B cell migration has been well demonstrated, but was also shown to be not the exclusive trigger [17], we were interested to find additional signals required in this process. Among the chemokine receptors, also *CCR7* and *CX3CR1* were differentially expressed. In the spleen and blood, *CCR7* expression remained consistently high, whereas in the bursa, *CCR7* expression significantly decreased from ED14 to ED16. CCR7 is known to guide lymphocyte migration to secondary lymphoid organs by responding to its ligands, CCL19 and CCL21 [30]. Since blocking of CCR7 and CXCR4 reduced the ability of B cells populating the lymph nodes in mice, both chemokine receptors seem essential in B cell homing of lymph nodes and B cells are triggered to adhere to either CCL21-expressing endothelial cells or CXCL12 expressed by stromal cells on high endothelial venules (HEVs) [31]. Moreover, it has been shown that CCR7 can modulate *CXCR4* expression and control CXCR4 responsiveness of B cells and bone marrow homing by inactivating CXCR4 [32]. Since chickens lack lymph nodes and the bursa is not considered as secondary lymphoid tissue, CCR7 may have a unique function in avian B cell immunology, distinct from its role in mammals. Our findings suggest that CCR7 play a role in directing chicken B cells to the bursa, in addition to CXCR4. The comparable expression levels of *CXCR4* and *CCR7* in the blood could indicate a coordinated process of both chemokines regulating each other.

B cells need to have specific cell surface molecules to colonize the bursa that allow them to pass through the ECM and bursal epithelium. It is suggested that B cells enter the bursa through HEVs, similar to lymphocyte diapedesis or transendothelial migration of leukocytes in mammals [33]. This extravasation process involves rolling of leukocytes along the endothelium, followed by integrin activation that enables firm adhesion and transmigration. Each step relies on specific receptors, creating a complex and coordinated interaction between homing receptors on leukocytes and cell adhesion molecules expressed on the counterpart [34, 35]. Tethering is mediated mainly by selectins, particularly through selectin L (SELL), which binds to its ligands expressed on HEVs [23, 35]. However, SELL has not been identified in chickens so far. Intriguingly, SELL ligands, along with ADAM17 - responsible for SELL’s cleavage and shedding - are annotated in the chicken genome and exhibit distinct expression patterns consistent with the DEGs identified in this study (Fig. 5D). However, there is currently no confirmation of SELL’s presence in chickens (Fig. S5), and its ligands may function in other pathways. Further investigation is needed to determine whether other selectin family members, such as SELE or SELP, may have replaced SELL’s functions or if selectins are not required for tethering of chicken leukocytes. In addition, while the surface glycosylation patterns of migrating lymphocytes and the role of (sialyl)-Lewis(x) in mediating adhesion could not be analyzed directly, the enzymes required for the synthesis of (sialyl)-Lewis(x) were identified (Fig. S3A). However, further research is needed to examine which specific fucosyl- and sialyltransferases are required to synthesize chicken (sialyl)-Lewis(x).

It has been shown that integrins are activated by chemoattractants; this facilitates migration and provides survival signals to cells during their passage through the ECM [35, 36]. An increase of *ITGAV*, *ITGB1*, *ITGB2*, *ITGA6* and *ITGB1BP1* expression in the blood on ED16, along with a decrease in the expression of integrins *ITGB1*, *ITGA6, ITGAV, ITGB2* and *ITGB4* in bursal B cells over time indicates that these integrins are essential during the migration process but become less critical once B cells have successfully entered the bursal mesenchyme. PECAM1, a member of the Ig superfamily, is expressed on both leukocytes and endothelial cells known to mediate leukocyte transmigration [37]. Since *PECAM1* expression significantly decreased in bursal B cells from ED14 to ED16, PECAM1 likely enables the physical movement of B cells across endothelial barriers, particularly during their transition from the bloodstream to the bursa. JAM3, another Ig superfamily member, is localized within endothelial tight junctions, primarily supporting cell adhesion through weak bonds, but does not directly adhere to mammalian leukocytes [38]. However, it was shown that using JAM3 antibodies diminished the migration of human JAM3-expressing B cells to secondary lymphoid organs [39], suggesting a more complex role in B cell extravasation that may vary between species. Elevated *JAM3* expression in the blood B cells at the latest time point and in the bursa at ED12 suggests that JAM3 might also be important in facilitating B cell transmigration in chickens, maybe through interaction with ITGB1 and ITGB2. Interestingly, while downregulation of *CCR7*, *CX3CR1*, integrins and CAMs in the bursa over time was observed, *CXCR4* expression remained unchanged. This pattern indicates that these signals are crucial for guiding B cells into the bursal mesenchyme, whereas CXCR4 is still needed for their subsequent colonization of follicle buds [16, 29].

The release of immature B cells from the bone marrow to the blood has been linked to S1PR1, whereas an interplay between CXCR4 and S1PR1 enabled the egress of plasma cells from the spleen [26, 40, 41]. The peak in *S1PR1* expression at ED14 coincides with heightened B cell exit from the spleen [42], indicating a potential role for S1PR1 also regulating chicken B cell egress. Further research is needed to determine whether higher S1P levels in the blood, relative to those in the tissue, create a gradient [24, 25] that also aids chicken B cells to exit the spleen.

The generation of JH-KO chickens, missing surface Ig, showed that the BCR itself is not required for B cells to enter the bursal mesenchyme [15]. However, it has been shown that CXCR4/CXCL12-driven B cell migration relies on BCR-mediated signaling to initiate this process [17]. Nevertheless, B cells must carry specific surface markers to distinguish them from other circulating immune cells, such as T-cells. Most likely, B cells selectively colonize the bursa rather than non-B cells leaving the bursa shortly after migrating into it. Therefore, DEGs associated with B cell activation and BCR signaling pathways were investigated (Tab. S4 and Fig. S4). Interestingly, several genes/proteins present in the BCR-mediated signaling pathway were also identified as differentially expressed in this study, suggesting that the BCR is likely involved in triggering the signaling cascade necessary for B cell migration. This signaling might be important for the exclusive immigration of B cells and could also contribute by activating pathways such as MAPK, PI3K-Akt and NF-κB which are essential for regulating the actin cytoskeleton, cell survival and B cell ontogeny [43]. Among others, these pathways were also identified as significantly enriched (Fig. 5B). Further research is needed to determine the underlying downstream signaling pathways and investigate whether these signals act in a BCR-dependent manner.

This study has identified promising candidates responsible to direct progenitor B cells to the bursa and colonization of the bursal mesenchyme in the chicken embryo. Our findings demonstrated that B cell migration is likely regulated by a coordinated process involving chemokines, integrins and cell adhesion molecules. We hypothesize that S1PR1 modulates B cell appearance in the blood, whereas guiding the B cell towards the bursa is facilitated by chemokine receptors CCR7 and CXCR4. Finally, cell adhesion molecules like PECAM1 and specific integrins mediate transmigration across endothelial barriers. This is a complex, multistep process involving several mediators essential for the recruitment and extravasation of progenitor B cells into the bursal mesenchyme during chicken embryonic development.

## Data Limitations and Perspectives

This section is not applicable to the current study.

## Material and Methods

### Animals

Chicken eggs from Lohmann-selected white Leghorn chickens were used for incubation at the Technical University of Munich Animal Research Center. Eggs were incubated at 37.8°C and 55% humidity and were automatically turned six times within 24 hours. All experiments were performed in accordance with the German Welfare Act and the European Union Normative for Care and Use of Experimental Animals.

### Isolation of PBMCs

Incubated chicken eggs were prepared for *in-ovo* blood collection as described before [44]. Depending on the embryós age, a maximum of 500-800µl blood was collected per embryo. Heparin-Natrium (50I.E./1ml RPMI, Ratiopharm GmbH, Ulm, Germany) was used to coat 1ml syringes and 27G needles, and the extracted blood per time point was combined. The pooled blood samples were diluted 1:1 with ice-cold 1xPBS. The cell suspension was layered onto histopaque-1077 (Sigma-Aldrich, Taufkirchen, Germany) in a 1:1 ratio, and density-gradient centrifugation was performed with 650xg for 12 min at RT.

### Isolation of the embryonic spleen and bursa

Embryonic spleens and bursae were processed in pools, with each pool consisting of two to five organs, depending on the age of the embryo. Spleens were homogenized by resuspending with a 1ml syringe in 1ml 1xPBS with 1% BSA. Single-cell suspensions of bursae were generated by passing them through a 100µm nylon Cell-Strainer with a 1ml syringe as a plunger. The pooled cell suspensions were diluted 1:1 with ice-cold 1xPBS, and lymphocytes were isolated using histopaque density-gradient centrifugation as described above.

### Flow-cytometry-based B cell sorting

Flow-cytometry and cell sorting were conducted as described before [45]. Briefly, PBMCs were washed with 1ml ice-cold 1xPBS at 222xg for 5 mins at 4°C. B cells were stained with anti-chicken Bu1-FITC clone AV20 (2.5 µg/ml) antibody (Southern Biotech, Alabama, United States), and live/dead staining was carried out with the 7-AAD-Viability Staining Solution (5 µg/ml) according to manufacturer instructions (BioLegend, San Diego, California, United States). Cell Sorting was performed using the CytoFlex SRT Benchtop Cell Sorter (Beckmann Coulter Life Sciences, Krefeld, Germany). B cells were directly sorted in 250-500µl BL+TG Lysis Buffer (ReliaPrep^TM^ RNA Cell Miniprep System, Promega, Walldorf, Germany). Data analysis was conducted with FlowJo^TM^10 v10.7.1 (BD, New Jersey, United States).

### RNA extraction

RNA isolation of sorted B cells followed the manufacturer’s instructions (ReliaPrep^TM^ RNA Cell Miniprep System, Promega, Walldorf, Germany). Total RNA was quantified using Nanodrop (ThermoFisherScientific, Schwerte, Germany), and RNA integrity was analyzed using the RNA Pico Chip on a 2100 Bioanalyzer (Agilent, Santa Clara, United States). Only B cell samples with a RIN > 8 were selected for RNA-sequencing.

### RNA-sequencing

Following the manufacturer’s instructions, the total RNA was utilized for library preparation with the SMART-Seq®-v4 PLUS Kit (Takara Bio Europe SAS, Saint-Germain-en-Laye, France). The quality of the obtained cDNA library was assessed using the Qubit®-dsDNA HS Assay Kit (ThermoFisherScientific, Schwerte, Germany) and the Agilent Bioanalyzer High-Sensitivity Kit (Agilent, Santa Clara, United States). Next-Generation-Sequencing (NGS) was performed by IMGM Laboratories GmbH (Martinsried, Germany) using the Illumina NovaSeq®-6000 SP v1.5 flowcell and chemistry (Illumina, San Diego, United States). Libraries were sequenced as single-end 1×100bp reads.

### Gene expression analysis

The sequence data have been submitted to the GenBank database under accession number PRJNA1190344. Examining raw sequencing reads involved quality control using FastQC v0.12.1 [46] and visualization through MultiQC v1.11 [47]. Read filtering and adapter trimming were executed with fastp v0.23.2 [48]. Alignment to the chicken genome *gallus gallus GCRg7w* (NCBI accession number: GCA_016700215.1) was conducted using HiSAT2 v2.2.1 [49] and gene count tables were generated using featureCounts v2.02.1 [50]. Genes with counts ≤ 10 in at least one B cell sample were excluded from further analysis. In total 8370 genes passed the criteria and were normalized using Median Ratio Normalization, batch-corrected with Combat, and pairwise comparisons performed for tissues and time points. Hierarchically clustering of the genes was performed with Euclidean distance and K-means clustering.

### Differential gene expression analysis

Differential gene expression analysis was performed with DeSeq2 v1.42.1 [51]. Genes were considered differentially expressed between the tissues and time points if p_adjust_ < 0.001 and fold-change > ⎢2⎢. The data analysis mentioned above was performed on a local Galaxy instance [52] and with DEBrowser v1.30.2 [53]. InteractiVenn [54] was used to visualize the number of shared DEGs across different tissues and time points. Gene expression profiles were generated, and Person-Correlation for Nearest-Neighbor analysis was performed with MORPHEUS [55]. The expression profiles of gene families or SELL interactors were visualized with GraphPad Prism v10.1.1 (GraphPad Software, Boston, Massachusetts, USA).

### Enrichment analysis

Gene Ontology (GO) analysis was carried out using Panther [56] with *gallus gallus* as the reference organism. PPI analysis was performed with the STRING App and functional enrichment using STRING Enrichment in Cytoscape v3.10.2 [57]. KEGG and Reactome pathways potentially involved in embryonic B cell migration were identified using DAVID [58] and visualized with the EnrichmentMap function in Cytoscape. To identify DEGs located on the cell surface, the gene set ‘cell surfac’ (GO ID: 0009986) was downloaded from MGI [59] and analyzed using RStudio. The KEGG pathway cell adhesion molecules (gga04514) and BCR signaling pathway (hsa04662) were retrieved and adapted from the KEGG PATHWAY Database [43].

### Statistical analysis

The statistical analysis was performed using GraphPad Prism v10.1.1 (GraphPad Software, Boston, Massachusetts, USA). Normally distributed data (as determined by Shapiro-Wilk test with a p-value < 0.05) were analyzed using one-way ANOVA with multiple comparisons. Statistical significance (p-values < 0.05) between the time points within each tissue is indicated in the figures.

## Abbreviations

BP: biological process
bp: base pair
BCR: B cell receptor
BSA: bovine serum albumin
CAMs: cell adhesion molecules
CC: cellular component
DEGs: differentially expressed genes
ECM: extracellular matrix
ED: embryonic day
GO: Gene Ontology
HEVs: high endothelial venules
HSCs: hematopoietic stem cells
Ig: immunoglobulin
MF: molecular function
NGS: Next-Generation Sequencing
PBMCs: peripheral blood mononuclear cells
PC: protein class
PCA: Principal Component analysis
PPI: protein-protein interaction
RNA: ribonucleic acid
S1P: sphingosine-1-phosphate
S1PR1: sphingosine-1-phosphate receptor 1
sIg: surface immunoglobulin

## Data availability statement

The data supporting the findings of this study can be obtained from the corresponding author upon reasonable request. The raw data for the transcriptome analysis were uploaded to GenBank under accession number PRJNA1190344.

## Ethics approval statement for human and/or animal studies

All animal work was conducted according to relevant national and international guidelines for the humane use of animals.

## Patient consent statement

Not applicable

## Permission to reproduce material from other sources

Not applicable

## Clinical trial registration

Not applicable

## Author contributions

MB, CC, TB and TvH performed the experiments. MB, TB and BS designed the experiments and wrote the manuscript. MB analyzed the data. BS supervised the project. DE performed the phylogenetic analysis. CW prepared the libraries and provided technical support for NGS. All authors read and approved the submitted version.

## Acknowledgments, including funding

This project was funded by the Deutsche Forschungsgemeinschaft (DFG, German Research Foundation) in the framework of the Research Unit ImmunoChick (FOR5130) project SCHU2446/6-1 (awarded to BS) and an Emmy-Noelther research fellowship (DFG SchU2446/3-1 awarded to BS). TB was supported by the Alexander von Humboldt Foundation during part of this study (NLD 1211322 HFST-P).

## Figures

All figures are included as a separate PDF file, following the submission guidelines. Each figure is accompanied by its corresponding legend in the main manuscript.

## Supporting Information

All supporting information’s are submitted as a separate PDF file.

