## Supporting Information for "Regulation of B cell migration to the Bursa of Fabricius by CCR7 and cell adhesion molecules in chicken embryonic development"

### Supp.1

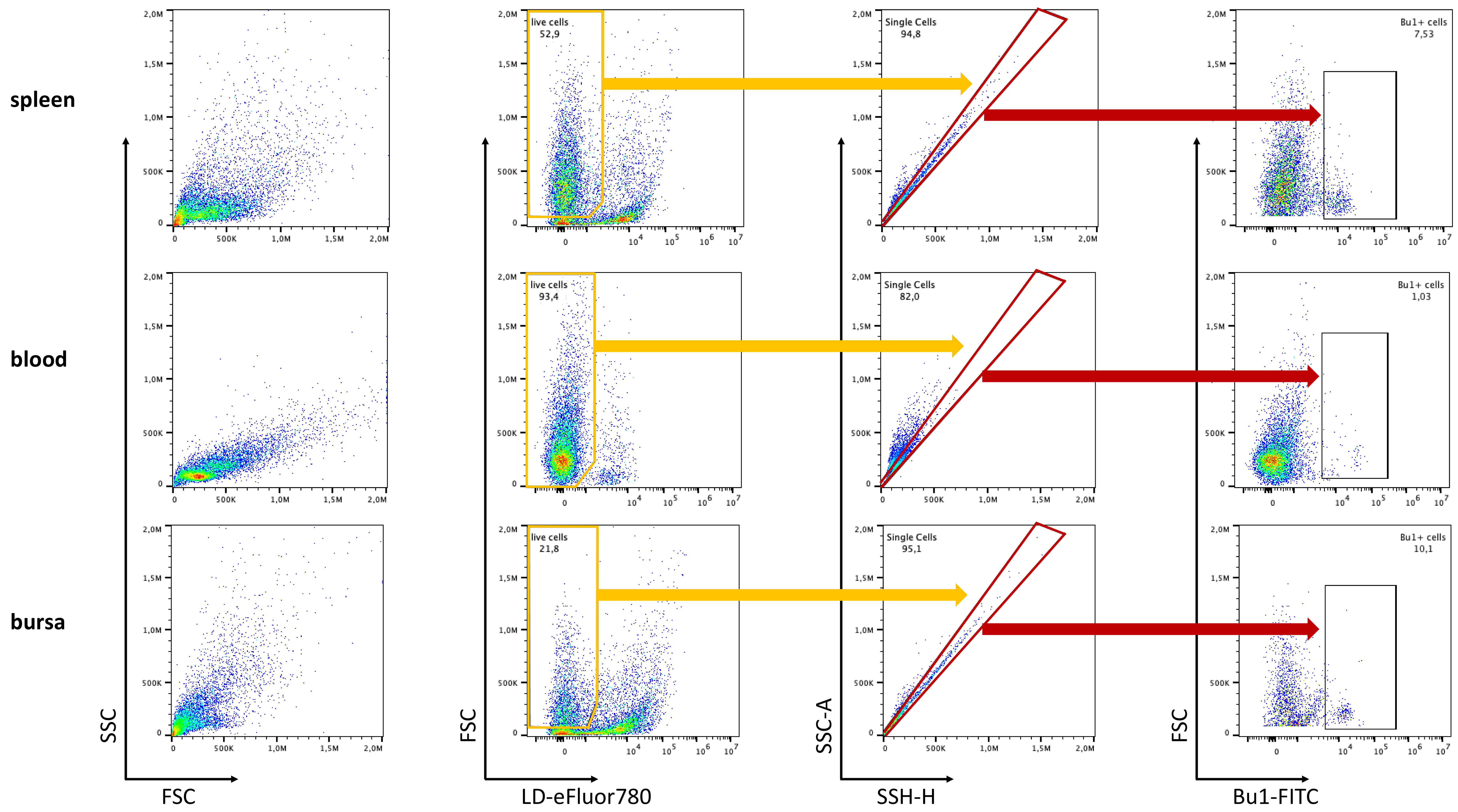


**Fig. S1:** Schematic representation of the flow-cytometry gating strategy for B cell percentages in the different tissues. Flow-cytometry analysis of B cells (Bu1-FITC$^{+}$) within single live cells (LD-eFluor780$^{-}$) in percentages from the spleen (top), blood (middle), and bursa (bottom).

Supp.2

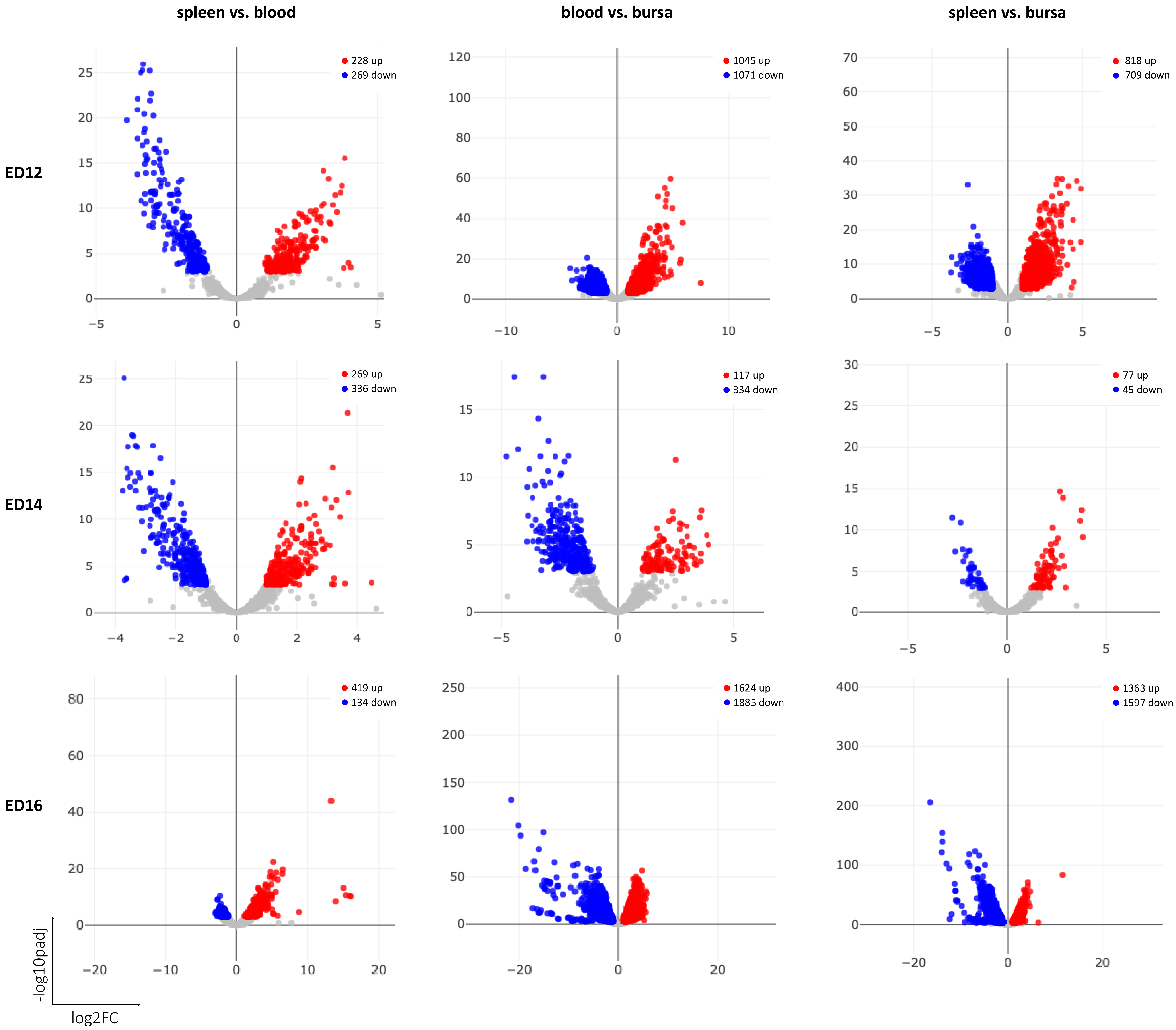


**Fig. S2:** Volcano plots of the up- and downregulated DEGs between the spleen vs. bursa (left), blood vs. bursa (middle), and spleen vs. bursa (right) on ED12 (top), ED14 (middle), and ED16 (bottom). The $log_2$(FC) reflects the mean fold-change (FC) in expression levels for each gene, and each dot represents one gene. DEGs are defined as $p_{adjust} < 0.001$, FC > |2|. DEGs with higher expression (i.e., higher in the spleen than in blood) are represented by red dots, while DEGs with lower expression (i.e., lower in the spleen than in blood) are denoted by blue dots. Genes with no substantial change in expression are indicated as gray dots.

Supp.3

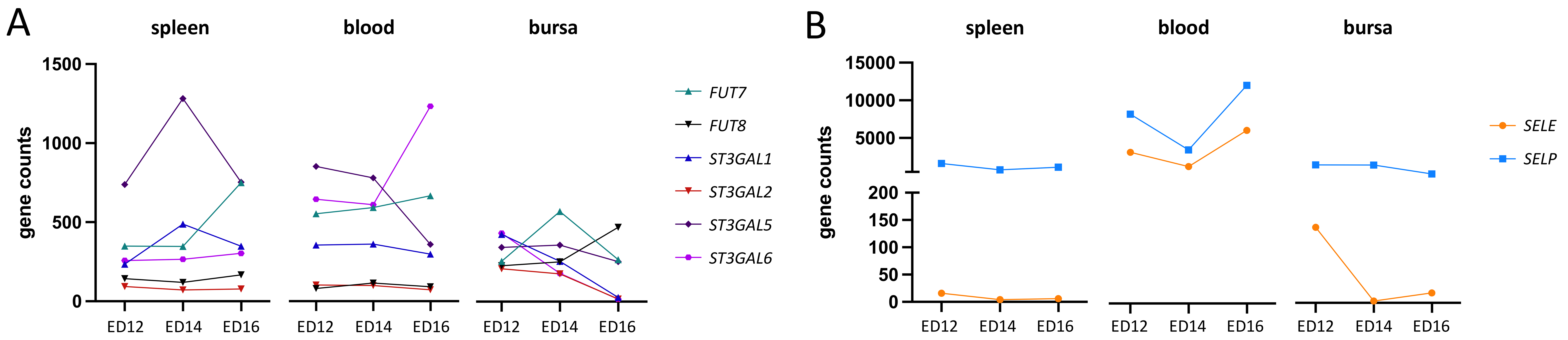


**Fig. S3:** Gene expression profiles of the (A) fructosyl- and sialyl transferases and (B) selectin E and P over time (ED12, ED14, and ED16) in the spleen (left), blood (middle), and bursa (right). Each symbol and color indicates one individual gene, data are cumulative from 3 experiments (*n*=3 per B cell sample and time-point) and represented as mean.

Supp.4

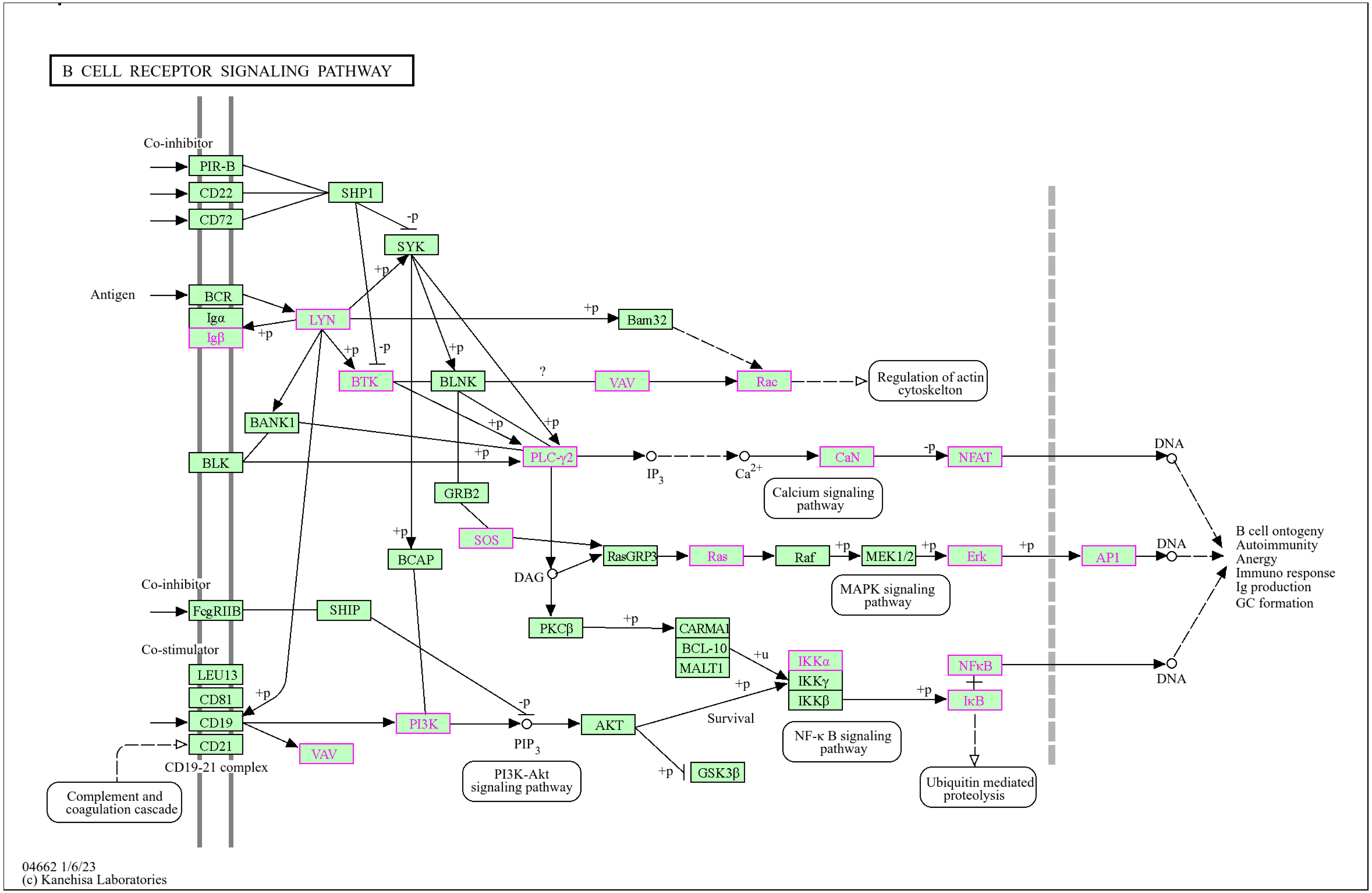


**Fig. S4:** KEGG pathway map for human B cell receptor signaling pathway. DEGs associated with the GO term B cell activation (Tab. S4) in the bursa at least at one time point comparison and mapped to the B cell receptor signaling pathway are marked in pink.

Supp.5

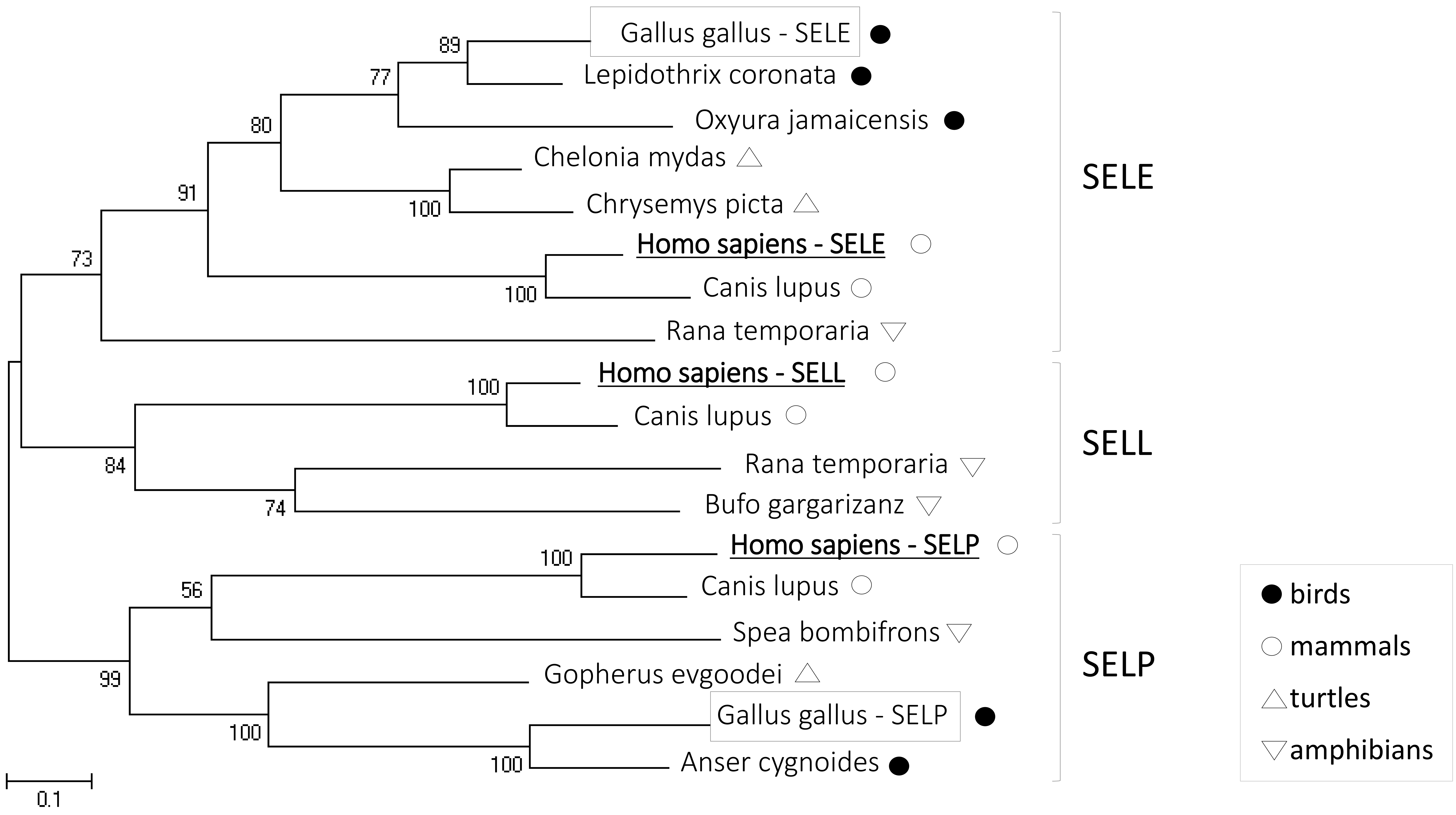


**Fig. S5:** Phylogenetic relationship of chicken selectins with other vertebrate orthologs. Both chicken selectins are highligted by grey boxes. Human E-selectin (SELE), L-selectin (SELL) and P-selectin (SELP) are shown in bold and underlined. Maximum likelihood tree was generated in MEGA6.06 software. Bootstrap support values (out of 100) are shown at each node. Scale bar shows number of amino acid substitutions per site. The following sequences were used to generate the phylogenetic tree: human SELL (accession number NP000646), human SELE (NP000441), human SELP (NP002996), *G. gallus* SELE (XP046754151), *G. gallus* SELP (XP422207), and selectin genes from various vertebrate species *Canis lupus* (XP025286208, XP025286205, XP025286209), *Rana temporaria* (XP040215647, XP040215645), *Bufo gargarizans* (XP044156813), *Spea bombifrons* (XP053326045), *Chelonia mydas* (XP037762768), *Gopherus evgoodei* (XP030427921), *Anser cygnoides* (X066857252), *Lepidothrix coronata* (XP017671135), *Oxyura jamaicensis* (XP035189260), *Chrysemys picta* (XP_065410664).

Supp.6

**Tab. S1:** Shared DEGs associated to the GO terms locomotion and cell adhesion molecules (n=37) between the spleen vs. blood, blood vs. bursa and spleen vs. bursa on ED12, ED14 and ED16

| **ED12** | | **ED14** | | **ED16** | |
|---|---|---|---|---|---|
| shared between all: spleen vs. blood, blood vs. bursa and spleen vs. bursa | NET1 | shared between all: spleen vs. blood, blood vs. bursa and spleen vs. bursa | | shared between all: spleen vs. blood, blood vs. bursa and spleen vs. bursa | PLXNC1<br>ITGB1<br>JAM3 |
| shared between spleen vs. blood and blood vs. bursa | | shared between spleen vs. blood and blood vs. bursa | ITGB2 | shared between spleen vs. blood and blood vs. bursa | BSG<br>RAC2<br>ITGAV<br>ITGA6 |
| shared between spleen vs. blood and spleen vs. bursa | ITGB1<br>ITGAV<br>MCAM | shared between spleen vs. blood and spleen vs. bursa | | shared between spleen vs. blood and spleen vs. bursa | CNTN1<br>CXCR4 |
| shared between blood vs. bursa and spleen vs. bursa | EPHB6<br>CYFIP1<br>TRIO<br>BSG<br>KIF5B<br>PIKFYVE<br>SEMA7A<br>RHOH<br>ELF1<br>PLXNA1<br>FAR2<br>ANTXR2<br>THSD7B<br>SGCE | shared between blood vs. bursa and spleen vs. bursa | | shared between blood vs. bursa and spleen vs. bursa | CCR7<br>NOTCH1<br>UNC5B<br>CX3CR1<br>CYFIP1<br>CYFIP2<br>SEMA4B<br>ATRN<br>MADCAM1<br>PCDH12<br>ITGB2 |

### Supp.7

**Tab. S2:** Protein-protein interaction (PPI) network analysis represents the associated GO term, FDR and p-value, number of background genes and genes included in the network

| term name | description | FDR value | # background genes | # genes in network | p-value |
|---|---|---|---|---|---|
| GO:0009986 | Cell surface | 1.68E-10 | 499 | 15 | $1.26E^{-13}$ |
| GO:0005886 | Plasma membrane | 3.2E-8 | 3837 | 28 | $4.79E^{-11}$ |
| GO:0031224 | Intrinsic component of membrane | 4.13E-8 | 4295 | 29 | $9.28E^{-11}$ |
| GO:0098636 | Protein complex involved in cell adhesion | 7.69E-8 | 32 | 6 | $2.31E^{-10}$ |

Supp.8

**Tab. S3:** List of genes identified in the gene set ´cell surface´ in the comparisons spleen vs. bursa and blood vs. bursa

| **cell surface** | | | |
|---|---|---|---|
| spleen vs. bursa | SELP | blood vs. bursa | FAS |
| | TSPAN32 | | PRNP |
| | TNFRSF10B | | FLT3 |
| | GOT2 | | HSP90AA1 |
| | C1QBP | | SCUBE1 |
| | HSP90AB1 | | MSN |
| | HMMR | | CKAP4 |
| | HSPD1 | | EPHB6 |
| | CD83 | | TNFRSF1A |
| | CLIC4 | | APP |
| | PRNP | | ITGB1 |
| | PDGFB | | LYPLA1 |
| | FCER1G | | HSP90AB1 |
| | CD8A | | CLU |
| | SCNN1B | | CTSB |
| | KCNH1 | | ANXA5 |
| | B2M | | ANXA2 |
| | ABCB11 | | MFGE8 |
| | BTN1A1 | | ADAMTS7 |
| | PECAM1 | | GOT2 |
| | VSIR | | LRRC8A |
| | CIITA | | TIMP2 |
| | IL15RA | | LBP |
| | TLR2 | | ANXA4 |
| | TGFB1 | | SLC9A1 |
| | NRROS | | JAM3 |
| | S1PR1 | | S100A10 |
| | BMPR2 | | IL2RG |
| | TIMP2 | | TLR2 |
| | CTNND1 | | KIT |
| | LYPLA1 | | CD44 |
| | VTCN1 | | CXCR4 |
| | GLRA1 | | NRROS |
| | ADAMTS7 | | HMMR |
| | MFGE8 | | P2RX7 |
| | CLSTN1 | | PECAM1 |
| | TNFRSF1A | | NTSR1 |
| | CD99L2 | | BTN1A1 |
| | LRRC8A | | |
| | ANXA2 | | |
| | APP | | |
| | ITGB1 | | |
| | CTSB | | |
| | SULF2 | | |
| | LBP | | |
| | JAM3 | | |
| | DCBLD2 | | |
| | HSP90AA1 | | |
| | ITGB2 | | |
| | LRP8 | | |
| | CTSZ | | |
| | HSPA8 | | |

**Tab. S4:** List of DEGs identified between the different time-points in the bursa associated with the GO term B cell activation

| **B cell activation** | | | | | |
|---|---|---|---|---|---|
| ED12 vs. ED14 | NFKB2<br>GRAPL<br>NFKB2<br>JUN<br>NFKBIA<br>NFKB1<br>HRAS<br>NFKBIA<br>PRKCD<br>PIK3CG<br>NFKB1<br>JUN<br>ITPR1 | ED14 vs. ED16 | CD79B<br>GRAPL<br>NFKBIA<br>CHUK<br>NFKBIA<br>PIK3CG<br>VAV3 | ED12 vs. ED16 | NFATC3<br>RAC1<br>PRKCD<br>MAPK8<br>JUN<br>CHUK<br>PPP3CA<br>SOS2<br>NFATC2<br>PIK3CG<br>PIK3CD<br>BTK<br>JUN<br>PLCG2<br>NFKB2<br>MAPK9<br>MAPK3<br>HRAS<br>LYN<br>BTK<br>LOC107050724<br>PTPRC<br>NFKB2 |
